# Single-cell analysis of skeletal muscle macrophages reveals age-associated functional subpopulations

**DOI:** 10.1101/2022.02.23.481581

**Authors:** Linda K. Krasniewski, Papiya Chakraborty, Chang-Yi Cui, Krystyna Mazan-Mamczarz, Christopher Dunn, Yulan Piao, Jinshui Fan, Changyou Shi, Tonya Wallace, Cuong Nguyen, Isabelle A. Rathbun, Rachel Munk, Dimitrios Tsitsipatis, Supriyo De, Payel Sen, Luigi Ferrucci, Myriam Gorospe

## Abstract

Tissue-resident macrophages represent a group of highly responsive innate immune cells that acquire diverse functions by polarizing towards distinct subgroups. The subgroups of macrophages that reside in skeletal muscle (SKM) and their changes during aging are poorly characterized. By single-cell transcriptomic analysis, we found that mouse SKM macrophages primarily comprise two large populations, “healing” LYVE1+ and “proinflammatory” LYVE1-macrophages. SKM macrophages were further classified into four functional subgroups based on the expression levels of another cell-surface marker, MHCII: LYVE1+/MHCII-lo (similar to alternatively activated M2), LYVE1-/MHCII-hi (similar to classically activated M1), and two new subgroups, LYVE1+/MHCII-hi and LYVE1-/MHCII-lo. Notably, the new subgroup LYVE1+/MHCII-hi had traits of both M2 and M1 macrophages, while the other new subgroup, LYVE1-/MHCII-lo, expressed high levels of mRNAs encoding cytotoxicity proteins. Flow cytometric analysis validated the presence of the four macrophage subgroups in SKM. In old SKM, LYVE1-macrophages were more abundant than LYVE1+ macrophages. Furthermore, complementary unsupervised classification revealed the emergence of specific macrophage subclusters expressing abundant proinflammatory markers, including *S100a8* and *S100a9* in aged SKM. In sum, our study has identified dynamically polarized mouse SKM macrophages and further uncovered the contribution of specific macrophage subpopulations to the proinflammatory status in old SKM.

## Introduction

Macrophages are heterogeneous innate immune cells (Shapouri-Moghaddam et al., 2018) that provide the first line of defense against pathogens but are also deeply involved in inflammation, dead-cell removal, wound healing, and tissue remodeling (Mills et al., 2014, Ross et al., 2021, Shapouri-Moghaddam et al., 2018). Macrophages adapt to individual tissues and acquire specific tissue-dependent functions (Wynn et al., 2013). Upon transplantation, tissue-resident macrophages quickly lose their original gene expression patterns and gain host organ markers (Lavin et al., 2014). The tissue environment was shown to determine tissue-specific protein production in macrophages and thereby establishes tissue-dependent expression patterns and functions (Gautier et al., 2012, Lavin et al., 2014). Hence, the function of macrophages should be studied in the context of their tissue of residence.

Macrophages play diverse functions in tissues by differentiating into specific functional subgroups, a process usually defined as macrophage polarization (Yao et al., 2019). Most macrophages are known to polarize to proinflammatory M1 or anti-inflammatory M2 subgroups (Martinez et al., 2008, Mills et al., 2000, Rath et al., 2014). While such dichotomy largely explains the strikingly different actions of macrophages commonly seen in many tissues, macrophages appear to be more functionally heterogeneous than simply M1 or M2. In fact, recent flow cytometry and single-cell studies have identified several new macrophage subgroups in arterial, lung interstitial, and adipose tissues (Chakarov et al., 2019, Jaitin et al., 2019, Lim et al., 2018, Schyns et al., 2019) with distinct tissue-dependent polarization status. Dissecting polarization in each tissue is thus critical to elucidate shared and tissue-dependent macrophage functions.

Skeletal muscle (SKM) contains large numbers of macrophages that play critical roles in injury repair and regeneration (Arnold et al., 2007, Tidball, 2017, Tidball, 2011). Macrophages assume different polarization to play distinct functions at different stages of repair after injury (Scala et al., 2021, Yang and Hu, 2018). In the absence of injury or infection, most macrophages residing in human and mouse SKM were shown to be CD206+ M2-like macrophages (Cui et al., 2019, Wang et al., 2015). However, the full range of macrophage subgroups and their age-related changes in SKM remain incompletely understood (Cui and Ferrucci, 2020).

To better understand the complexity of macrophage polarization status and their changes with aging in mouse SKM, we carried out single-cell transcriptomic analysis. We present evidence that SKM macrophages can be divided into two large populations by a surface marker, LYVE1, and can be further classified into four functional subgroups by introducing MHCII as an additional surface marker. We further show that complementary, unsupervised classification identified additional smaller macrophage subgroups, and that inflammatory biomarkers including the mRNAs that encode S100A8/9 and FABP4/5 were significantly upregulated in specific subsets in old SKM. Our findings reveal dynamically polarized functional subpopulations of mouse SKM macrophages, including changes towards a proinflammatory status with aging.

## Results

### Isolation of macrophages from mouse SKM and single-cell RNA-sequencing

To isolate macrophages from SKM, we collected all muscles from hind limbs, including quadriceps, gastrocnemius, tibialis, and soleus, from C57BL/6JN mice, combined and minced them into small cubes, and isolated mononuclear cells by digesting them with enzymes including collagenase and proteases (Krasniewski et al., 2021, Liu et al., 2015) (Fig. 1A). To identify macrophage-rich fraction from the mononuclear cell preparation, we carried out flow cytometric analysis based on the presence of CD45, a pan-leukocyte marker, and CD11b, a pan-myeloid lineage marker. As shown in our previous report, CD11b+ cells clearly separated from the rest of the mononuclear cell population, and virtually all CD11b+ cells were CD45+ (Krasniewski et al., 2021).

**Fig. 1:**
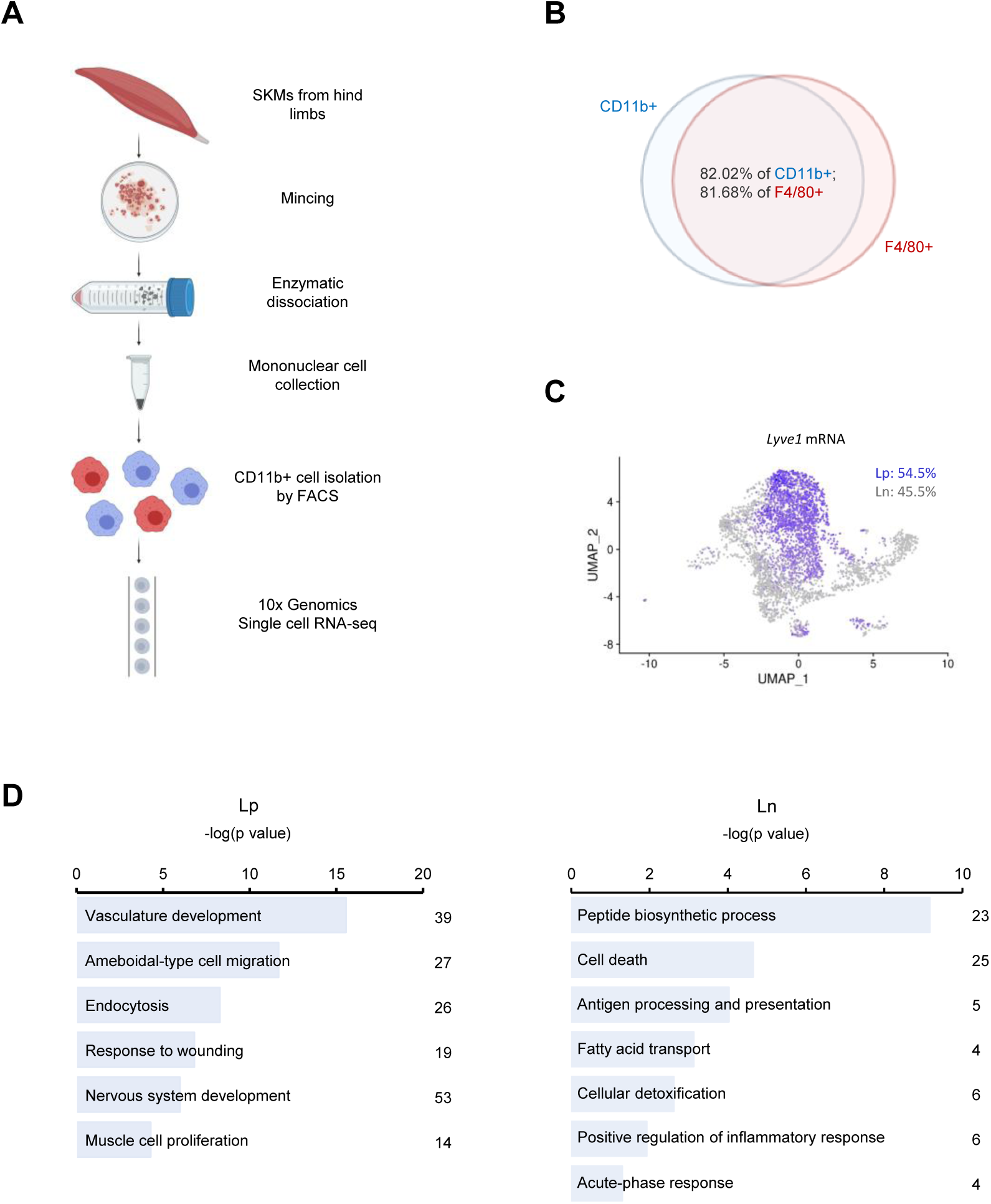
Macrophage isolation from mouse SKM and single-cell RNA-seq analysis. (A) Workflow of mononuclear cell collection from mouse SKM, CD11b+ cell isolation by FACS, and single-cell RNA-seq analysis using the 10x Genomics platform. (B) Cells isolated from mouse SKM that were CD11b+ and F4/80+ (another marker of mouse macrophages in single-cell RNA-seq analysis). (C) Separation of SKM macrophages into groups according to LYVE1 levels; ∼54.5% of macrophages were LYVE1+, and ∼45.5% were LYVE1-. (D) GO annotation of the distinct functions of Lp and Ln macrophages, with ‘healing’ features for LYVE1+ macrophages and antigen processing/presentation and inflammation features in LYVE1-macrophages. Lp, LYVE1+. Ln, LYVE1-.

For single-cell RNA-sequencing (RNA-seq) analysis, we collected CD11b+ cells from 3 young [3 months old (3 m.o.)] male mice as biological triplicates by fluorescence-activated cell sorting (FACS). From each mouse, 5,000 to 10,000 CD11b+ isolated cells were subjected to single-cell library preparation using the 3’ gene expression pipeline from 10X Genomics followed by RNA-seq analysis. We successfully obtained sequences from 3,000-5,000 single cells from each mouse, and a mean of ∼80,000 RNA-seq reads per cell corresponding to a median of >2000 genes per cell (Materials and Methods; GEO identifier GSE195507; token for reviewers is etifyqaexrcrjmh). As anticipated, sequencing analysis showed that more than 80% of *CD11b+* cells were also positive for *F4/80* (ADGRE1), another popular marker for mouse macrophages (Fig. 1B). We analyzed CD11b+ and F4/80+ double-positive cells as SKM macrophages in this study. Very few cells were positive for *Ly6G* mRNA or *SiglecF* mRNA (specific markers for neutrophils and eosinophils, respectively; Fig. S1A), indicating that there was minimal contamination from these cells in our macrophage population.

Initially we attempted to subgroup our SKM macrophages by well-accepted traditional polarization markers: CD206 (MRC1), CD86 and CD80. CD206 is a widely used marker of M2 macrophages, whereas CD80 and CD86 are known as M1 markers (Mantovani et al., 2002, Stein et al., 1992). However, our single-cell RNA-seq data showed that *Cd206* and *Cd86* mRNAs were broadly expressed in more than 80% of macrophages, and *Cd80* mRNA was expressed only in a small population (Fig. S1B). Furthermore, most macrophages expressed *Cd206* and *Cd86* mRNAs simultaneously (Fig. S1B). These observations indicated that traditional polarization markers are not ideal to classify SKM macrophages at the transcriptomic level.

### SKM macrophages comprise two large functional populations, LYVE1+ and LYVE1-

We therefore turned to other candidate markers to subgroup SKM macrophages. Expression of LYVE1 was recently used to subgroup arterial and lung interstitial macrophages (Chakarov (Chakarov et al., 2019, Lim et al., 2018). LYVE1 status divided SKM macrophages into two large, similarly sized groups, *Lyve1*+ (Lp, 54.5%) and LYVE1-(Ln, 45.5%) (Fig. 1C). To gain insight into their function, we performed Gene Ontology (GO) enrichment analysis (g:Profiler). Consistent with the macrophage characteristics, both Lp and Ln populations expressed mRNAs encoding proteins with typical macrophage functions, including “immune system process”, “defense response”, “regulation of cell adhesion”, and “response to stress”. However, Lp macrophages were further enriched in mRNAs encoding proteins important for “healing”, including angiogenesis, cell migration, endocytosis, wound healing, and muscle cell proliferation (Fig. 1D, Lp), while Ln macrophages showed elevated expression of mRNAs encoding ribosomal proteins, cell death-related proteins, antigen-processing and antigen-presenting proteins, proinflammatory proteins, antioxidants, and fatty acid transporters (Fig. 1D, Ln).

We found that Lp macrophages displayed an M2-like transcriptomic program, which included mRNAs encoding proteins with roles in “vascular development”, “wound repair”, and “endocytosis” (Fig. 2A, *left*) (Buchacher et al., 2015, Stein et al., 1992). Interestingly, transcripts encoding proangiogenic proteins (*Ang, Stab1, Egr1*, and *Fgfr1* mRNAs) as well as transcripts encoding anti-angiogenic proteins (*Cfh* and *Hspb1* mRNAs) were upregulated in Lp macrophages. Similarly, transcripts encoding proteins important for wound healing (*Igf1, Nrp1, Hbegf*, and *Gas6* mRNAs) were elevated in Lp macrophages (Fig. 2A, *left*); mRNAs encoding endocytosis-related members of the CD209 family (including *Cd209f, Cd209d, Cd209g*, and *Cd209b* mRNAs) as well as *Cd163* and *Cd206*/*Mrc1* mRNAs were also highly expressed in Lp macrophages. Additional mRNAs, such as *Fcna* and *Timd4* mRNAs, were almost exclusively expressed in the Lp macrophages and hence might be good candidate markers for this population (Fig. S2A). In contrast, Ln macrophages expressed high levels of many mRNAs encoding ribosomal proteins (Fig. S2B), as well as cell death-related proteins (*Plac8, Bcl2a1b*, and *Btg1* mRNAs), proinflammatory proteins (*Il1b* and *Adam8* mRNAs), antioxidants (*Gsr, Prdx5*, and *Hp* mRNAs), fatty acid transporters (*Fabp5* and *Fabp4* mRNAs), and antigen-presenting proteins (*H2-Eb1, H2-Ab1*, and *Cd74* mRNAs) (Fig. 2A, *right*). The full list of mRNAs differentially abundant in Lp and Ln is listed in Table S1.

**Fig. 2:**
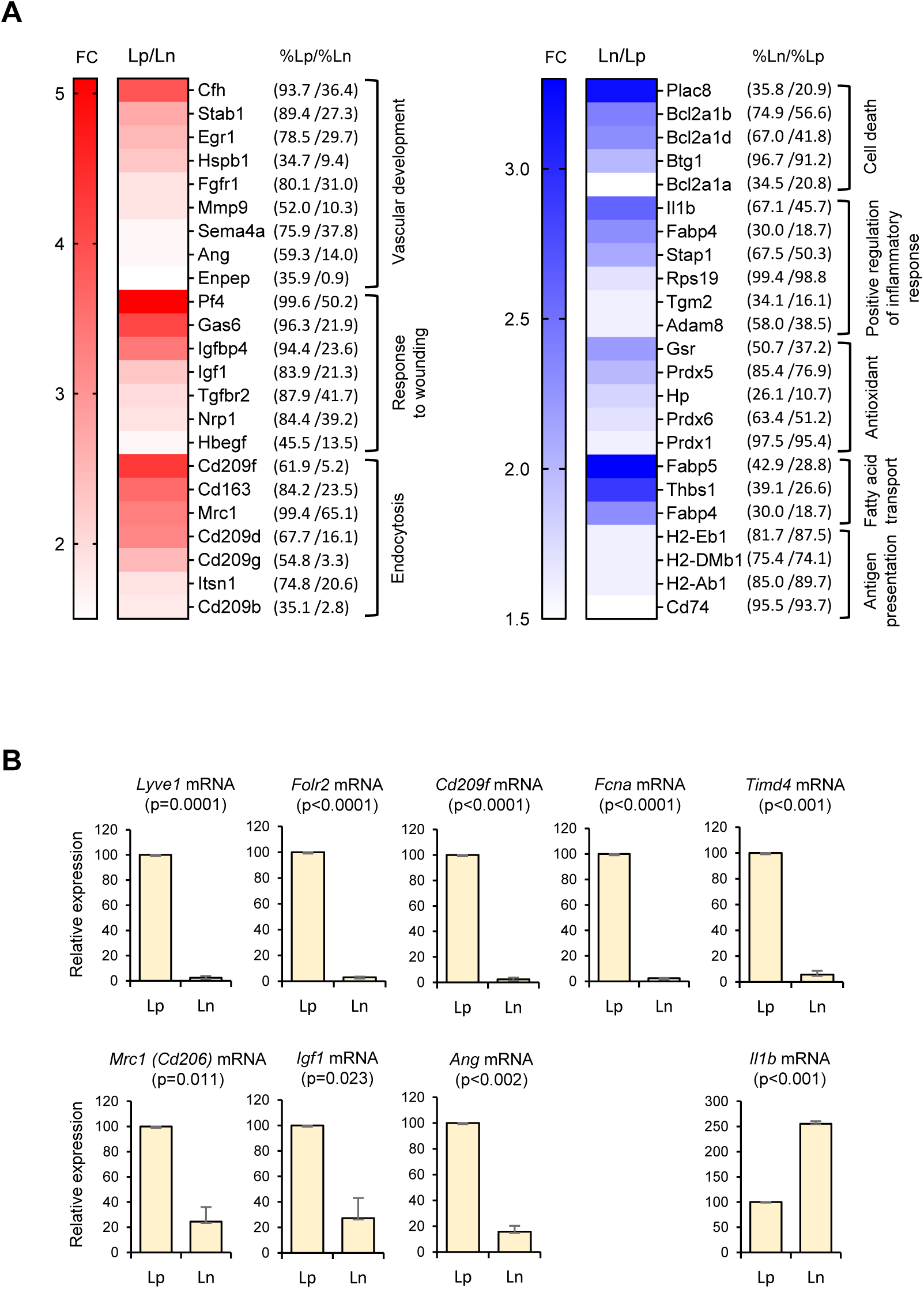
Functional clusters of genes differentially expressed in Lp and Ln macrophages following single-cell RNA-seq analysis. (A) Left panel, genes highly expressed in functional clusters of Lp macrophages. Expression fold changes (Lp/Ln) and percentages of positive cells in Lp and Ln subpopulations (%Lp/%Ln) are shown. Right panel, genes highly expressed in Ln macrophages. Expression fold changes (Ln/Lp) and percentages of positive cells in Lp and Ln subpopulations (%Ln/%Lp) are shown. (B) Validation of select mRNAs differentially abundant as identified in panel (A). Lp and Ln macrophages were isolated by FACS analysis from 3 male mice, 3 months old (m.o.), and mRNAs elevated in Lp macrophages (bottom left and top), and mRNAs predominantly elevated in Ln macrophages (bottom right) were quantified by RT-qPCR analysis. Data were normalized to the levels of *Gapdh* mRNA, also measured by RT-qPCR analysis. Data represent the means and S.D. from two different sorts for each group.

To validate the differences in gene expression programs, we separated each population (Lp and Ln macrophages) by FACS analysis, isolated total RNA from each population, and performed reverse transcription (RT) followed by real-time quantitative (q)PCR analysis (RT-qPCR). Lp (CD45+/CD11b+/F4/80+/LYVE1+) and Ln (CD45+/CD11b+/F4/80+/LYVE1-) macrophages comprised ∼45% and ∼55% of total macrophages (CD45+/CD11b+/F4/80+), respectively (see below). RT-qPCR analysis showed that *Lyve1, Folr2, CD209f, Fcna, Timd4* mRNAs were almost exclusively expressed in Lp (Fig. 2B, top graphs, n=2 biological replicates). By contrast, *Cd206, Igf1*, and *Ang* mRNAs were expressed in both Lp and Ln macrophages, but at much higher levels in Lp, while *Il1b* mRNA was significantly elevated in the Ln population (Fig. 2B, bottom graphs). The RT-qPCR results (Fig. 2B) were consistent with the single-cell transcriptomic analysis (Fig. 2A, Fig. S1C, and Table S1), indicating that LYVE1 is an effective marker for subgrouping mouse SKM macrophages.

### MHCII proteins further classify SKM macrophages into four functional subgroups

To delineate the diverse functions of macrophages at finer resolution, we sought to further subdivide SKM macrophages. MHCII has been used to classify lung interstitial and arterial macrophages along with LYVE1 (Chakarov et al., 2019, Lim et al., 2018). We found that MHCII mRNAs (encoding H2-Ab1, H2-Eb1) divided SKM macrophages into two groups, MHCII-high (Hh) and MHCII-low (Hl) in single-cell profiling analysis. The relative abundance of LYVE1 and MHCII allowed the classification of SKM macrophages into four subgroups: LYVE1+/MHCII-low (LpHl), LYVE1+/MHCII-high (LpHh), LYVE1-/MHCII-high (LnHh), and LYVE1-/MHC-low (LnHl) (Fig. 3A). Among them, LpHh and LnHh were the largest subgroups, comprising 38.5% and 35.5% of all macrophages, respectively (Fig. 3A), while LpHl and LnHl comprised 15.4% and 10.6%, respectively. The gene expression patterns from the single-cell analysis (heat map, Fig. 3B) were clearly distinct across the four subgroups. Those mRNAs that were expressed >1.5-fold higher in a given subgroup relative to the other 3 subgroups, p<0.01, and were expressed in >25% of macrophages in that subgroup are shown in Table S2.

**Fig. 3:**
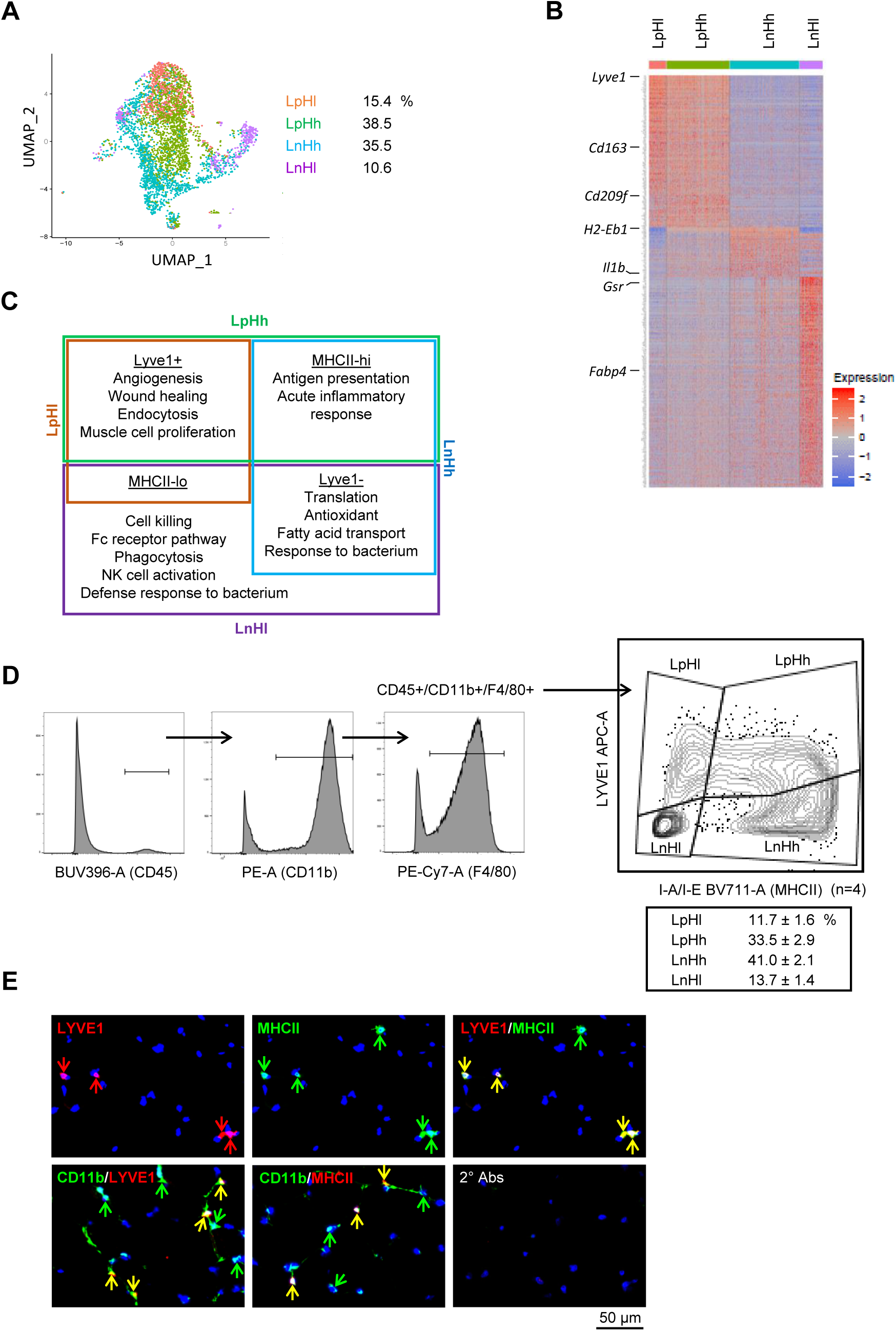
Classification of mouse SKM macrophages into four functional subgroups. (A) Subclassification of mouse SKM macrophages based on LYVE1 and MHCII levels: LpHl, LpHh, LnHh, and LnHl. UMAP analysis shows the distribution and size of the four subgroups. (B) Heat map analysis of the single-cell RNA-seq data depicting the gene expression patterns of the four macrophage subgroups. (C) GO annotation of the functions of each subgroup. Brown box, LpHl; green box, LpHh; blue box, LnHh; purple box, LnHl. (D) Flow cytometric analysis of the four subgroups in SKM. CD45+/CD11b+/F4/80+ macrophages (left 3 panels show gating) were further classified by LYVE1 and MHCII (right). LpHl, LnHh, and LnHl subgroups formed clear cell clusters, while LpHh spanned LpHl and LnHh. Note: the sizes of each subgroup by flow cytometric analysis were similar to those seen with single-cell RNA-seq analysis. Gating was based on FMO (fluorescence minus one) controls for each experiment. (E) Immunofluorescence analysis of the presence of LpHh macrophages in mouse SKM. *Top*, LYVE1+, MHCII+, and LYVE1+/MHCII+ double-positive cells in endomysium and perimysium areas of mouse SKM. *Bottom*, colocalization of LYVE1 (left) and MHCII (middle) with CD11b, a macrophage marker. *Bottom* right, secondary antibodies only.

The functional annotations of the genes showing differential expression in each subgroup were identified. LpHl macrophages (*brown box*, Fig. 3C) were associated with angiogenesis and wound healing, similar to M2 macrophages (Krzyszczyk et al., 2018). LnHh macrophages (*blue box*, Fig. 3C) showed higher antigen-presentation activity, acute inflammatory response, active translation and antioxidant functions, and were more like M1 macrophages (Mills, 2015). LpHh macrophages (*green box*, Fig. 3C) were more complex; they largely shared LpHl (M2-like) functions, including angiogenesis and wound healing, but also shared LnHh (M1-like) functions, such as antigen processing, antigen presentation, and acute inflammatory response. By contrast, LnHl (*purple box*, Fig. 3C) showed more cytotoxicity, in addition to strong translation and antioxidant functions. Notably, among the four subgroups, the LpHh and LnHl subgroups have not been reported in SKM (Wang et al., 2020), and LnHl macrophages were not reported in any other tissues so far (Chakarov et al., 2019, Lim et al., 2018). Thus, in addition to M2-like (LpHl) and M1-like (LnHh) subgroups, single-cell profiling analysis revealed two new functional subgroups, LpHh and LnHl, in young, resting mouse SKM. Table S3 includes full lists of genes differentially expressed in each subgroup.

We further analyzed if the macrophage subgroups identified from single-cell RNA-seq could be validated by cell-surface protein markers. As expected, flow cytometric analysis using antibodies that recognized LYVE1 or MHCII divided CD45+/CD11b+/F4/80+ SKM macrophages into four subgroups, LpHl, LpHh, LnHh and LnHl (Fig. 3D, n=4 biological replicates). Notably, the LpHl, LnHh and LnHl subgroups showed clear clusters of cells, but LpHh macrophages spread across LpHl and LnHh (Fig. 3D). The sizes of each subgroup identified by flow cytometry were comparable to the sizes of the subgroups identified by single-cell transcriptomics (compare the numbers in Fig. 3A and 3D).

To validate if these macrophage subgroups are present in mouse SKM, we performed immunofluorescence staining analysis (Fig. 3E). LYVE1+ cells (*red*) and MHCII+ cells (*green*) were found in the endomysium and perimysium regions. Importantly, many LYVE1+ cells were also MHCII+ (double-positive, LpHh) in mouse SKM (Fig. 3E, yellow arrows, *top right*), consistent with the flow cytometric and the single-cell transcriptomic analyses. By tyramide signal amplification (TSA) staining of CD11b with detection of LYVE1 and MHCII, we confirmed that LYVE1+ and MHCII+ cells were CD11b+ (Fig. 3E, *bottom*). Immunohistology thus confirmed that the new LpHh subgroup is a macrophage subpopulation found constitutively in mouse SKM.

### Macrophage subgroups show distinctive phagocytic capacities

To gain insight into functional differences among the four subgroups, we assessed their phagocytic capacity, a fundamental function of macrophages, by a flow cytometry-based method that measures the uptake of labeled particles (pHrodo Red *E. coli* Bioparticle assay, Materials and Methods). As anticipated, all macrophage subgroups were strongly phagocytic (Fig. 4A), with 90.9% of LpHl, 94.1% of LpHh, 85.8% of LnHh, and 49.1% of LnHl macrophages actively phagocytizing *E. coli* particles at 37°C; in control incubations, <3% of macrophages were active in each group at 4°C (Fig. 4B, n=3 biological replicates). The number of active phagocytic macrophages in the LnHl subgroup was lower than in the other three subgroups (Fig. 4B, p<0.01), but those that were active showed strikingly greater phagocytic capacity than the other three subgroups, as measured by geometric mean fluorescence intensity (gMFI) (Fig. 4C, n=3). The significance of the distinct behavior of this subgroup of SKM macrophages is unclear at present.

**Fig. 4:**
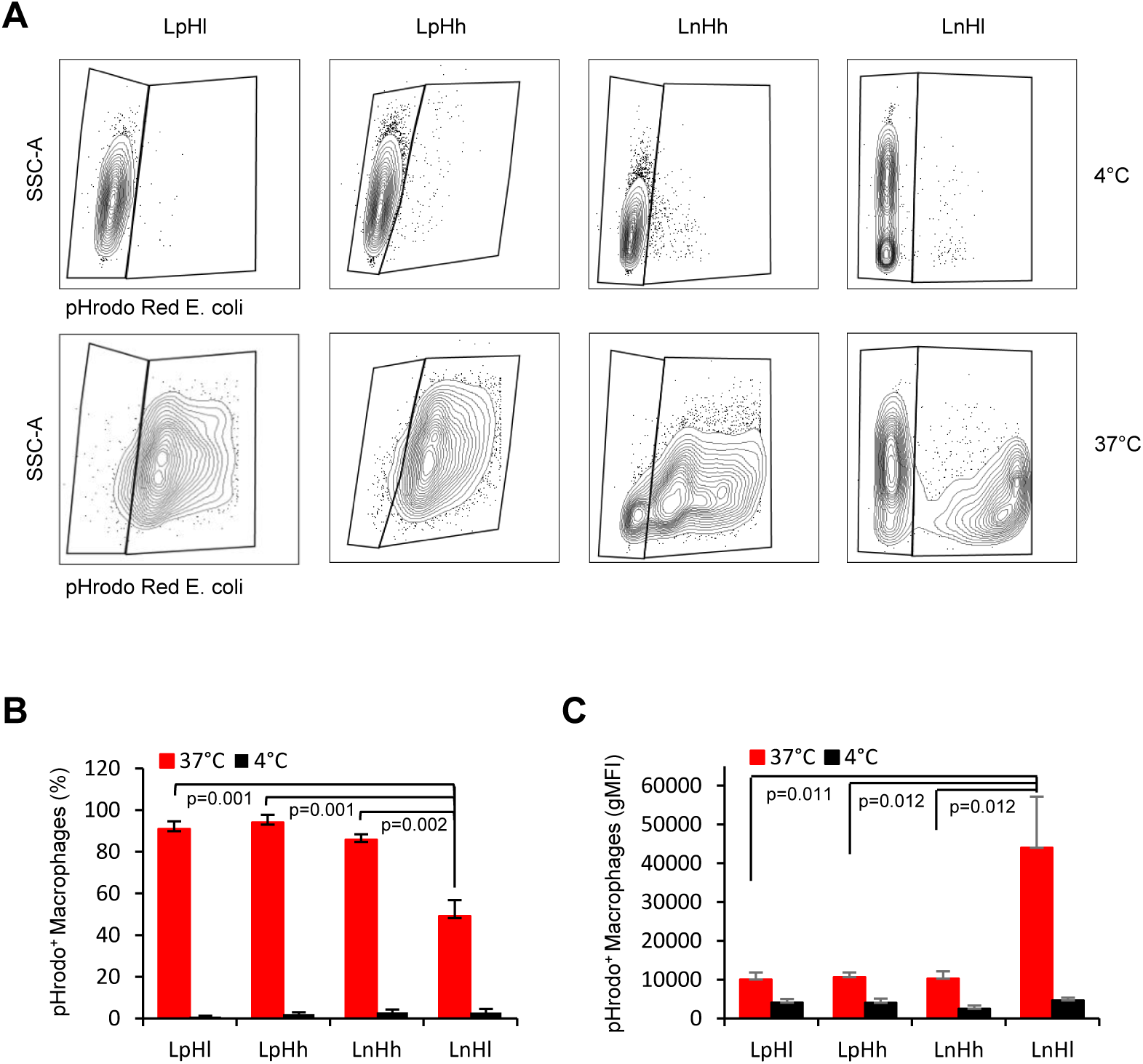
Analysis of the phagocytotic capacities of each macrophage subgroup. (A) Phagocytic activity was measured for mouse SKM macrophages at 4 °C (control, low phagocytosis) and 37 °C (active phagocytosis, right boxes). Gating was established using FMO controls for each experiment. (B) Quantification of the macrophages showing active phagocytosis in the four subgroups. (C) Quantification of phagocytic activity using geometric Mean Fluorescence Intensity. Data in (A) are representative of three experiments. Data in (b,c) represent the means and S.D. from three different sorts.

### Enrichment of proinflammatory, LYVE1-negative macrophages in aged SKM

We next analyzed how aging influenced the relative abundance of the four SKM macrophage subgroups. Single-cell RNA-seq analysis of old (23-m.o.) SKM macrophages (n=3 biological replicates) was compared with young (3-m.o.) SKM macrophages (Fig. 3A) (Materials and Methods). The results indicate that all four SKM macrophage subgroups are found in both age groups (Fig. 5A compared to Fig. 3A), but the percentage of LpHl and LpHh macrophages is slightly lower in the old SKM, while the percentage of LnHh and LnHl macrophages is higher in old mice (Fig. 5A, *right*). Flow cytometric analysis confirmed the same trend (Fig. S3A, n=4 biological replicates), as LpHh macrophages were lower and LnHh and LnHl macrophages were significantly higher in old mice (Fig. S3B, *top*). When considering a single marker, both the reduction in Lp and the rise in Ln were statistically significant (Fig. S3B, *bottom*). Thus, both single-cell transcriptomics and flow cytometry revealed that “healing” LYVE1+ macrophages decreased and “proinflammatory” LYVE1-macrophages increased in number in old mice.

**Fig. 5:**
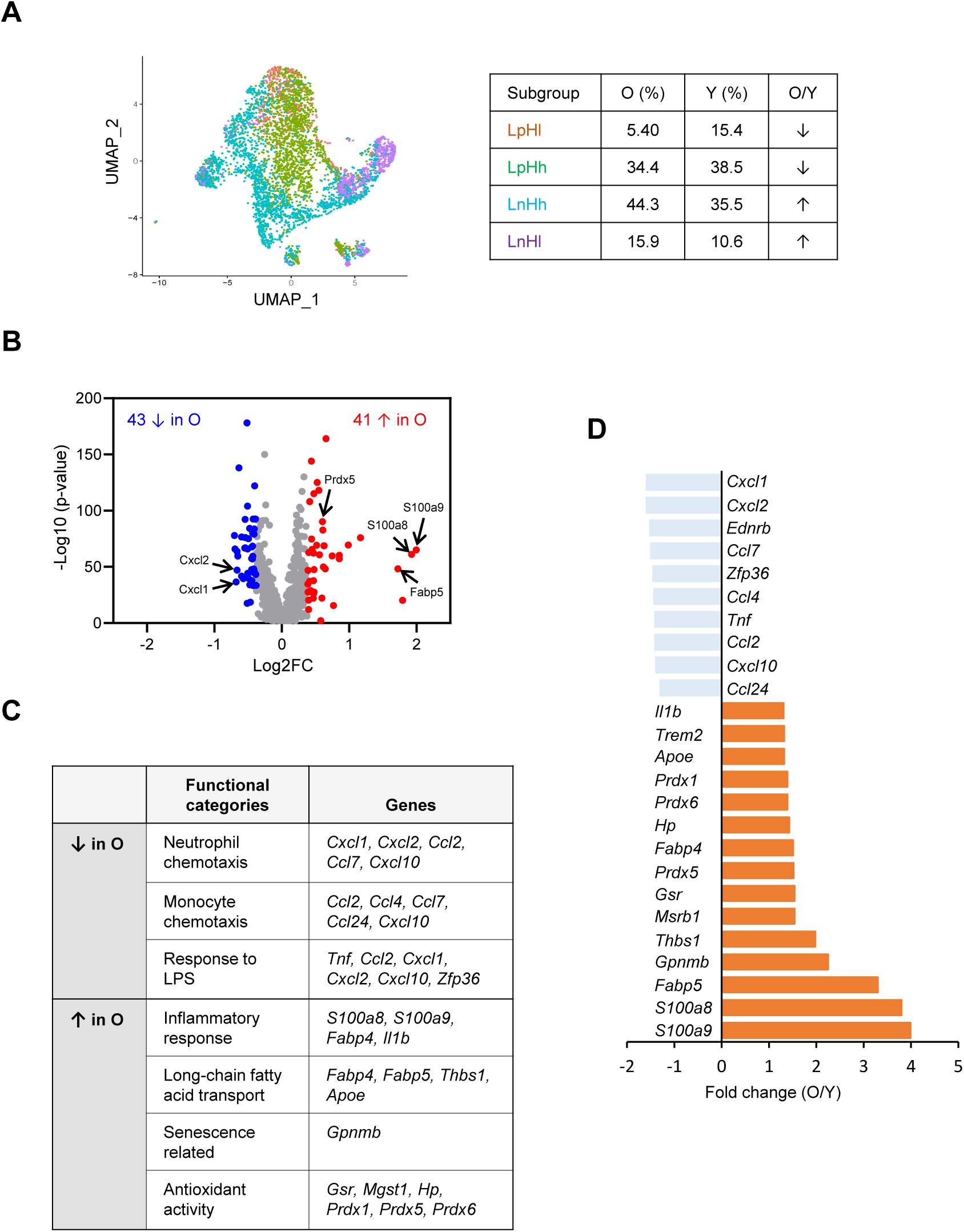
Analysis of gene expression programs in SKM macrophages from young and old mice. *Top*, UMAP representation of single-cell RNA-seq analysis from 23 m.o. old mice. *Bottom*, Relative proportions of LpHl, LpHh, LnHh, and LnHl in SKM from old (23 m.o.) and young (3 m.o.). By scRNA-seq analysis, a total of 84 genes were differentially expressed between old and young SKM. Arrows indicate upregulated (red) or downregulated (blue) mRNAs in old SKM macrophages. (C) GO annotation depicting the functional categories that were upregulated and downregulated in the old SKM macrophages relative to young SKM macrophages. (D) Fold changes in the abundance of select mRNAs (O/Y), as determined from the scRNA-seq analysis.

We further analyzed genes differentially expressed in macrophages from young and aged mouse SKM. We first compared gene expression programs without subgrouping. 41 mRNAs were elevated and 43 were reduced in macrophages from old SKM (Fig. 5B) following the criteria of mRNAs expressed in >10% of total macrophages in young or old, p < 0.01, and fold change > 1.3. The number of differentially abundant mRNAs was rather small, likely reflecting the healthy status of the SKM in this cohort, but the differences were informative. GO annotation showed that mRNAs encoding proteins involved in chemotaxis of neutrophils (e.g., *Cxcl1* and *Cxcl2* mRNAs) (Girbl et al., 2018) and monocytes (e.g., *Ccl2* and *Ccl7* mRNAs) (Deshmane et al., 2009), and the response to LPS (e.g., *Tnf, Cxcl10*, and *Zfp36* mRNAs) were downregulated in macrophages from old SKM (Fig. 5c,d). By contrast, mRNAs encoding proinflammatory proteins and biomarkers (e.g., *S100a8, S100a9*, and *Il1b* mRNAs) (Wang et al., 2018), long-chain fatty acid transporters (*Fabp4* and *Fabp5* mRNAs) (Babaev et al., 2011, Furuhashi et al., 2007), a candidate senescent-cell membrane marker (*Gpnmb* mRNA) (Suda et al., 2021), and antioxidant enzymes (e.g., *Gsr, Hp, Prdx1, Prdx5*, and *Prdx6* mRNAs) were elevated in old SKM macrophages (Fig. 5C,D). The full list of differentially expressed genes is shown in Table S4.

All four macrophage subgroups displayed differentially expressed mRNAs (Fig. S3C), suggesting that SKM macrophages were widely affected with age, although some mRNAs were altered only in certain subgroups. For example, *S100a9* mRNA was upregulated in all four subgroups in aged SKM. *S100a8, Cxcl1*, and *Cxcl2* mRNAs were differentially abundant in three subgroups, while *Il1b* mRNA was elevated, whereas *Ccl2* and *Ccl7* mRNAs were reduced only in the LnHl subgroup. Among the four subgroups, LnHl showed the largest number of mRNAs differentially expressed in young relative to old (Table S4). Overall, gene expression data suggested that anti-pathogen functions declined and proinflammatory functions were elevated in macrophages from old SKM.

### Unsupervised classification identified small macrophage clusters altered in old SKM

The presence of cell surface markers LYVE1 and MHCII (Chakarov et al., 2019, Lim et al., 2018) effectively subgrouped SKM macrophages through *supervised* classification. To complement this analysis, we performed *unsupervised* classification by gene expression patterns only. Unsupervised clustering provided ten clusters (Cls 0-9) of macrophages from young and old SKM using FindClusters resolution of 0.3 (Fig. 6A). Each cluster showed distinct expression patterns (Fig. 6B), and macrophages in the bigger clusters overlapped with those identified after supervised subgrouping. For example, Cl0 and Cl1 largely corresponded to LpHl and LpHh macrophages, respectively, while Cl2 and Cl4 macrophages overlapped with LnHh, and Cl3 with LnHl (compare Fig. 6A with Fig 3A and 5A). Among the larger clusters (Cls 0-4), Cl3 showed a big change in the old, similar to “supervised” LnHl (Fig. 6A and Fig. S3C).

**Fig. 6:**
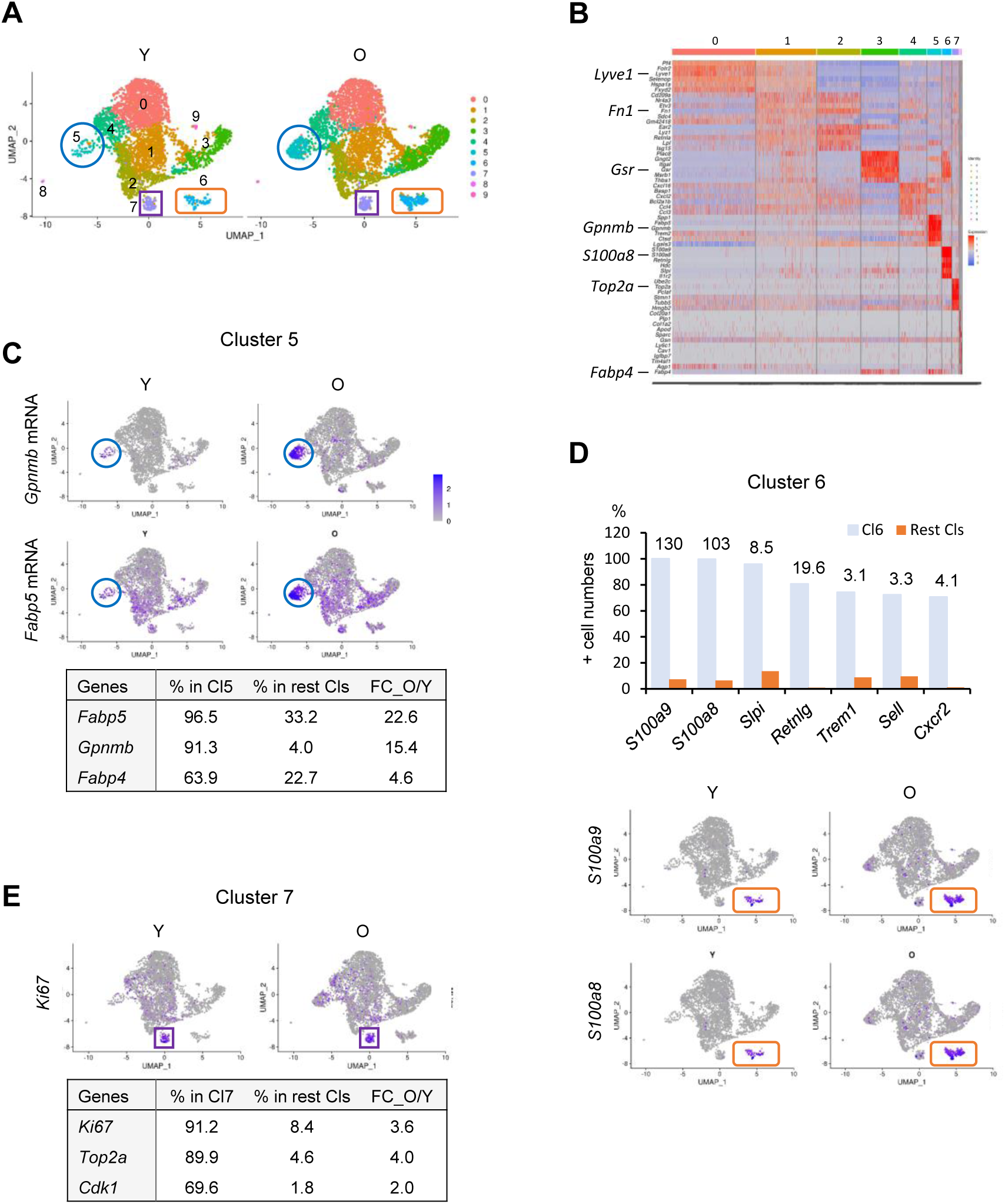
Identification of additional SKM macrophage clusters from scRNA-seq analysis followed by unsupervised classification. (A) Unsupervised clustering (UMAP analysis) revealed 10 macrophage clusters in young and old SKM; small clusters Cl5 (blue) and Cl6 (brown) showed significantly increased numbers of macrophages in the old. (B) Heat map analysis of gene expression patterns after unsupervised classification of mouse SKM macrophages illustrating the top mRNAs upregulated in each cluster. (C) *Top*, UMAPs analysis of *Gpnmb* and *Fabp5* mRNAs, elevated in old SKM macrophages in Cl5. *Bottom*, percentages of positive macrophages in Cl5 (% in Cl5) and in the combined other clusters (% in rest Cls), and expression fold changes between the old and young (FC_O/Y) for select mRNAs. (D) *Top*, graph representing the relative numbers of cells expressing chemoattractant mRNAs in O/Y SKMs; numbers above the bars represent expression fold changes in the abundance of such mRNAs between the old and young (O/Y) SKMs. Note: *S100a8* and *S100a9* mRNAs showed >100-fold elevation in the old relative to young SKMs in Cl6. *Bottom*, UMAP analysis of *S100a8* and *S100a9*, almost exclusively present in Cl6. (E) *Top*, similar numbers of *Ki67-*expressing SKM macrophages in young and old in Cl7; *bottom*, mRNAs encoding cell cycle-related factors (*Ki67, Top2a*, and *Cdk1* mRNAs), were almost exclusively expressed in Cl7 relative to all clusters (rest Cls) and were elevated in the old SKM.

Notably, smaller clusters like Cl5 and Cl6, which were mixed populations in the supervised subgrouping (Fig. 3A, 5A), showed striking differences in abundance between young and old SKM (Fig. 6A, blue circles and brown rectangles). GO annotation revealed that mRNAs encoding a putative senescence marker (*Gpnmb* mRNA) and proteins implicated in fatty acid transport and inflammation (e.g., *Fabp5* and *Fabp4* mRNAs) were predominantly expressed in Cl5 and were significantly more abundant in old SKM macrophages (Fig. 6C; full gene lists in Table S5).

By contrast, several mRNAs in Cl6, including *S100a8* and *S100a9* mRNAs, were almost exclusively expressed in this cluster and were strikingly elevated in old SKM (Fig. 6D; Table S5 for full gene list). Many of them encoded proteins involved in leukocyte chemotaxis (Fig. 6D). In particular, mRNAs encoding proinflammatory biomarkers S100A8 and S100A9 were upregulated more than 100-fold in macrophages from old SKM in this cluster (Fig. 6D, *top*, numbers above the bars), further supporting the proinflammatory status of old SKM.

While Cl7 did not show significant changes in cell numbers between young and old (Fig. 6A, purple square), macrophages in this cluster expressed high levels of mRNAs encoding cell cycle-related proteins (*Ki67, Top2a*, and *Cdk1* mRNAs) (Fig. 6E; full list in Table S5), suggesting that Cl7 may represent a subpopulation of proliferating macrophages (Wang et al., 2020). Unexpectedly, these mRNAs were more highly expressed in the old macrophages (Fig. 6E, bottom); while *Ki67* mRNA was expressed in only ∼10% of total SKM macrophages, it was expressed in 91.2% of Cl7 macrophages. *Cdk1* and *Top2a* mRNAs were expressed in an even lower number of macrophages (data not shown). Thus, unsupervised classification of single-cell RNA-seq data identified smaller macrophage subpopulations that may play key functions, complementing the major subpopulations identified through supervised grouping.

## Discussion

Heterogeneity and functional versatility are critical characteristics of macrophages. Derived from embryonic and/or adult hematopoietic system (Cox et al., 2021), macrophages adapt their gene expression profile to the tissues in which they reside and play diverse functions by polarizing to different subgroups. In this study, we identified functional subgroups of mouse SKM macrophages by single-cell transcriptomic analysis. We found that SKM macrophages can be subdivided into populations that express or lack LYVE1 on their plasma membrane, and can be further divided into four functional subgroups by the levels of cell-surface MHCII proteins. These four subgroups covered well-known M2-like and M1-like macrophages, and two additional new subgroups that were confirmed by flow cytometry and immunohistology, while unsupervised classification after single-cell transcriptomic analysis further revealed additional functional clusters, some of which are differentially represented and/or acquire a pro-inflammatory expression profile in older compared to younger muscle. Thus, our study has characterized the diverse populations of macrophages in resting mouse SKM.

It is worth noting that the traditional polarization markers CD206 and CD86 (Mantovani et al., 2002, Stein et al., 1992) were widely and simultaneously expressed in most mouse SKM macrophages. We and others observed similarly wide expression of CD206 in mouse and human SKM macrophages by immunohistology (Cui et al., 2019, Wang et al., 2015) and flow cytometric analysis showed simultaneous expression of CD206 and CD86 in most human SKM macrophages (Kosmac et al., 2018). These observations suggest that for SKM macrophages, CD206, CD86 and CD80 may not properly represent polarization.

As alternatives, LYVE1 and MHCII were recently used to subgroup arterial and lung interstitial macrophages (Chakarov et al., 2019, Lim et al., 2018). Our study supports the notion that LYVE1 and MHCII are effective markers to delineate the polarization status of mouse SKM macrophages. Notably, the same markers have revealed differentially polarized macrophage subgroups in other tissues (Chakarov et al., 2019, Lim et al., 2018). Among four subgroups found in SKM, M2-like LpHl and M1-like LnHh subgroups were detected in both artery and lung, while the LpHh subgroup was found in artery (Lim et al., 2018), but not lung (Chakarov et al., 2019). By contrast, the LnHl subgroup was not reported in any other tissues so far. These observations support the notion of tissue-dependent polarization, and identify a population that appears to polarize selectively in SKM and not in artery or lung.

Among the four subgroups, the new LpHh subgroup showed both M1- and M2-like gene expression patterns and functional capabilities (Fig. 3C, D). We hypothesize that this subgroup may have distinct functions or may have the potential to shift to M2-like LpHl or M1-like LnHh subgroups depending on surrounding conditions. The gene expression heat map showed that LpHh expresses features of both LpHl and LnHh, but these patterns are not prominent (Fig. 3B). In flow cytometric analysis, LpHh macrophages spanned two clear clusters, LpHl and LnHh (Fig. 3D), possibly suggesting that LpHh macrophages represent an intermediate stage, even if they stand alone as an independent population (Fig. 3E). The function of LpHh macrophages with respect to LpHl and LnHh macrophages requires further study.

By contrast, the new LnHl subpopulation, which clearly separated from the other three subgroups by flow cytometric analysis (Fig. 3D), was predicted to have a more distinct “killing” capacity and may be directly implicated in innate immunity. In phagocytosis assays, the LnHl subgroup showed fewer active macrophages (Fig. 4B), but those that were active had strikingly stronger phagocytic capacity compared to the other three subgroups (Fig. 4C), further highlighting a distinct behavior by this specific subgroup. Genes known to be highly expressed in circulating monocytes, such as that encoding *Ly6c* mRNA (Wolf et al., 2019), was expressed in <3% of LnHl and the other subgroups, while *Cd115* (*Csf1r*) and *Ccr2* mRNAs were abundantly expressed in all subgroups (Table S2, and data not shown). CD11C, a dendric cell (DC) marker (Singh-Jasuja et al., 2013), and CD49 and CD122, candidate markers for lymphoid lineage NK cells (Nabekura and Lanier, 2016), were not detected in LnHl or the other subgroups (Table S2). These data strengthen the view that LnHl macrophages are distinct from circulating monocytes, or DC and NK cells. Additional studies are also needed to functionally characterize each macrophage subgroup, especially the two new subgroups LpHh and LnHl, in SKM.

Our study further revealed aging-related expression changes in macrophages in SKM. Overall, *healing* macrophages were less abundant, and *proinflammatory* macrophages were more abundant in aged SKM (Fig. 5A, Fig. S3A). Consistent with these observations, *S100a8* and *S100a9* mRNAs, encoding proinflammatory biomarkers, were significantly elevated in macrophages from aged SKM. Unlike neutrophils, macrophages were reported to express S100A8 and S100A9 at low levels in the absence of stimulation (Hessian et al., 1993, Wang et al., 2018). Often forming heterodimers, S100A8 and S100A9 serve as biomarkers for the diagnosis and therapeutic responses in inflammatory diseases, like inflammatory arthritis and inflammatory bowel disease, while blocking their activity resulted in reduced inflammation in mouse models (Wang et al., 2018). *S100a8* and *S100a9* mRNAs were in very low abundance in macrophages from young SKM but were strikingly more abundant in old SKM (Figs. 5D, 6D). The levels of *Fabp4, Fabp5*, and *Il1b* mRNAs, encoding additional proinflammatory proteins, were also upregulated in macrophages from old SKM (Figs. 5D, 6D). Macrophage-derived FABP4 and FABP5 were shown to promote a proinflammatory state in the vasculature during atherosclerosis development (Babaev et al., 2011, Furuhashi et al., 2007, Makowski et al., 2001). Increased expression of the above proteins in macrophages supports the proinflammatory status in old SKM. We propose that the expression levels of S100A8 and S100A9 in macrophages can be essential indicators of the inflammatory status of SKM, and possibly other tissues (Wang et al., 2018). We also found increased expression of mRNAs encoding antioxidant enzymes in old SKM macrophages, possibly reactive to elevated ROS in aged SKM (Jackson and McArdle, 2011).

By contrast, the mRNAs encoding several neutrophil and monocyte/macrophage chemoattractants (Deshmane et al., 2009, Girbl et al., 2018) were expressed in lower amounts by old SKM macrophages (Fig. 5C, D). In pathological conditions, like injury or infection, neutrophils are the earliest effector cells to infiltrate into the injury site followed by monocytes/macrophages (Forcina et al., 2020). At the same time, it is well known that in old SKM, injury repair and regeneration are slower, perhaps because leukocyte infiltration at early stages is delayed and CCAAT enhancer-binding protein beta (CEBPB) in macrophage polarization toward regeneration after muscle injury may be lower (Blackwell et al., 2015). Thus, the reduced production of chemoattractants in macrophages may contribute to the delay of repair in older SKM.

Finally, unsupervised classification identified additional smaller macrophage clusters (Fig. 6), which complemented the supervised classification. Many mRNAs defining the smaller subclusters in unsupervised classification were expressed in <25% of the macrophage populations in each of the four subgroups defined by supervised classification, and thus did not appear to be significant. Unsupervised clustering identified smaller functional subgroups including Cl5 and Cl6, which showed robust changes during aging (Fig. 6C, D). However, specific surface markers remain to be identified for each of the smaller subclusters. Notably, Cl7 contained proliferating macrophages that were reported in SKM recently (Wang et al., 2020). Many mRNAs encoding proliferation proteins, including KI67, TOP2A, and CDK1, were significantly elevated in old SKM macrophages (Fig. 6E). In line with this finding, macrophages were shown to proliferate by inflammatory cytokines secreted by senescent cells (Covarrubias et al., 2020). However, the biological significance of the proliferation-related genes expressed in this cluster of old SKM macrophages requires further analysis.

Aging impacts essentially all tissues and organs. Intrinsic and extrinsic factors, including DNA damage, endoplasmic reticulum stress, mitochondrial dysfunction, and systemic inflammatory environment in aged individuals inevitably affect the characteristics of macrophages (van Beek et al., 2019). A recent study suggested that macrophages from old SKM contribute to axonal degeneration and demyelination in the neuromuscular junction, and depletion of macrophages led to increased muscle endurance (Yuan et al., 2018). We propose that the age-associated SKM macrophage gene expression patterns identified here represent an important step towards elucidating how macrophage subpopulations influence the pathophysiology of old SKM.

## Materials and methods

### Collection of SKMs from young and aged C57BL/6JN mice

All animal study protocols were approved by the NIA Institutional Review Board (Animal Care and Use Committee). Young (Y, 3 months old, 3 m.o.) and aged (O, 22-24 m.o.) male inbred C57BL/6JN mice were purchased from the NIA aged rodent colony (https://ros.nia.nih.gov/). The mice were sacrificed, and all hind limb muscles, including quadriceps, hamstring, gastrocnemius, soleus, and tibialis anterior muscles were harvested as explained. Collected samples were directly used for mononuclear cell isolation or frozen in isopentane chilled by liquid nitrogen and stored at -80°C for immunohistology.

### Mononuclear cell isolation from SKM for flow cytometric analysis and single cell RNA-seq

Tendons, blood vessels, and fat tissues were removed under a dissection microscope. Muscle tissues were finely chopped and minced using dissection scissors to form a slurry. For single-cell RNA-seq analysis, we isolated mononuclear cells with Miltenyi’s Skeletal muscle dissociation kit (#130-098-305) with GentleMACS Octo Dissociator (#130-096-427), as described previously (Krasniewski et al, 2021). For further flow cytometric analysis, we also used an established method (Liu et al., 2015) with slight modifications. Briefly, the muscle slurry was digested with 1000 U/mL Collagenase type II (Gibco, Cat# 17101015) in 10 mL of complete Ham’s F-10 medium (Lonza, Cat# BE02-014F) for 70 min with 70 rpm agitation at 37 °C. Partially digested muscles were washed in complete Ham’s F-10 medium and centrifuged at 400 rcf speed for 5 min and cell pellet with 8 mL of the remaining suspension (pellet 1) was collected; 42 mL of the supernatant was collected in two tubes (21 mL each) that were filled up to 50 mL with Ham’s F10 media, and centrifuged again at 500 rcf for 8 min and the pellet (pellet 2) was collected. Pellet 1 was subjected to a second round of digestion in 1 mL of 1000 U/mL Collagenase type II and 1 mL of 11 U/mL Dispase II (Thermofisher, Cat# 17105041) along with the 8 mL of the remaining cell suspension, for 20 min with 70 r.p.m. agitation, at 37 °C. Digested tissues were aspirated and ejected slowly through 10 mL syringe with 20-gauge needle followed by washing in complete Ham’s F10 media at 400 rcf for 5 min. The supernatant was collected and centrifuged again at 500 rcf for 8 min and the pellet obtained (pellet 3) was pooled with the pellet 2 above. The suspension of pellet 2+3 was filtered through 40-μm cell strainer (Fisher scientific, Cat # 22363547), followed by final wash in complete Ham’s F10 medium. Cell pellets were resuspended in 1 mL complete Ham’s F10 medium. Cell counting was performed using trypan blue (Invitrogen, Cat# T10282) at a 1:1 ratio in CountlessTM cell counting chamber slides (Invitrogen, Cat# C10228) using CountlessTM II FL Automated Cell Counter (Invitrogen).

### Flow cytometric analysis and FACS

Flow cytometric analysis and CD11b+ cell isolation by FACS were described previously (Krasniewski et al, 2021). For flow cytometric validation studies, RT-qPCR (reverse transcription followed by quantitative polymerase chain reaction) analysis, and phagocytosis assays, mononuclear cell suspensions were incubated with BD Horizon™ Fixable Viability Stain 780 (FVS780, BDBiosciences, Cat# 565388, Dilution: 1:4000) in PBS (Ca+ and Mg+ free, Thermofisher) for 30 min at 4 °C in the dark. Fc receptors were blocked using TruStain FcX™ (anti-mouse CD16/32) Antibody (Biolegend, Cat# 101320, Clone: 93, Dilution 1:1000) for 5 min at 4°C in FACS staining buffer (1% BSA and 10 mM EDTA in Miltenyi’s Auto MACS Rinsing Solution). For macrophage sorting, mononuclear cells were further stained in FACS staining buffer for 40 min at 4 °C in the dark, with fluorochrome conjugated antibodies specific to mouse as indicated: BUV395 Rat Anti-Mouse CD45 (BDBiosciences, Cat# 564279, Clone: 30-F11, Dilution: 1:100), PE anti-mouse/human CD11b Antibody (Biolegend, Cat# 101208, Clone: M1/70, Dilution: 1:100), PE/Cyanine7 anti-mouse F4/80 Antibody (Biolegend, Cat# 123114, Clone: BM8, Dilution: 1:40), Brilliant Violet 711™ anti-mouse I-A/I-E Antibody (Biolegend, Cat# 107643, Clone: M5/114.15.2, Dilution: 1:40), APC Rat Anti-Mouse Lyve1 Antibody (Thermofisher, Cat# 50-0443-82, Clone: ALY7, Dilution: 1:20). Cells were fixed using BD Cytofix Fixation buffer (BD Biosciences, Cat# 554655) for 20 min on ice in the dark for analysis (but not for sorting). Compensation matrices were created using single color controls prepared using COMPtrol Kit, Goat anti-Mouse Ig (H&L) coated particles, with negative and high in separate vials (Spherotech, Cat# CMIgP-30-2K), combining one drop from each vial in equal ratio. The cells were acquired on a BD FACSAria™ Fusion (BD Biosciences) instrument and analyzed with Flowjo software (Tree Star, Inc).

### Macrophage single cell RNA-sequencing with Chromium Controller (10x Genomics)

Macrophages isolated from three 3 m.o. and three 23 m.o. C57BL/6JN male mice (biological triplicates) were stained with CD11b antibody, and isolated by FACS analysis. Single-cell libraries were prepared with 10x Genomics Chromium Single Cell 3’ Reagent Kits v3 (10x Genomics Cat# PN-1000092) with Chip B (10x Genomics, Cat# PN-1000073) following the manufacturer’s protocol with Chromium Controller (10x Genomics). Briefly, 5,000-10,000 single macrophages were used for GEM (Gel Bead-in-Emulsion) generation. The cDNAs were then synthesized and their qualities assessed on the Agilent Bioanalyzer with High Sensitivity DNA kit (Agilent Cat# 5067-4626). cDNAs were then used for library preparation and the quality of the final libraries assessed on the Agilent Bioanalyzer with DNA 1000 kit (Agilent, Cat# 5067-1504). The libraries were sequenced with Illumina Nova-sequencer with a depth of 300-400 million reads per sample. RNA-seq data are deposited in GEO with identifier GSE195507; token for reviewers is etifyqaexrcrjmh.

### Single-cell RNA sequencing data analysis

Single-cell RNA-seq samples were demultiplexed and mapped to the mm10 mouse reference genome using the Cell Ranger software version 3.0.2 (10X Genomics). Further analysis of the matrices of read counts obtained was carried out in R with the Seurat package, version 4.0.4 (Hao et al., 2021), using default parameters in all functions, unless specified otherwise. To exclude empty droplets, poor-quality cells and potential doublets from downstream analysis, quality control filtering was applied for each sample, which removed cells containing more than 7.5% mitochondrial genes, cells expressing <300 or >7,000 transcripts, and below 500 or above 60,000 counts. Genes that were detected in less than 10 cells were eliminated from the analysis. Cells expressing *Itgam* (*Cd11b*) and *Adgre1* (F4/80) mRNAs, two key macrophage markers, were subjected to further analyses. Each sample was normalized with the LogNormalize method, and the top 2,000 variable genes were used for integrated analysis, following scaling and Principal Component Analysis (PCA). To visualize and explore cell clusters in two-dimensional space, Uniform Manifold Approximation and Projection (UMAP) was performed using the first 30 principal components determined by the ElbowPlot method.

For *supervised* cluster analysis, the macrophage dataset was divided into four cell subgroups based on the log-normalized expression values of Lyve1 and H2-Ab1 (MHCII) mRNAs, as follows: LpHl (Lyve1 > 0 and H2-Ab1 < 2), LpHh (Lyve1 > 0 and H2-Ab1 ≥ 2), LnHh (Lyve1 ≤ 0 and H2-Ab1 > 2), and LnHl (Lyve1 ≤ 0 and H2-Ab1 ≥ 2). *Unsupervised* cell clustering was carried out by using FindNeighbors function and setting FindClusters resolution to 0.3, which distinguished 10 clusters of cells with distinct gene expression profiles. mRNAs that were expressed in at least 25% of cells per cluster were further considered for differential gene expression analysis among clusters. mRNAs were defined as differentially expressed if they had an absolute fold change >1.5 and adjusted p-value < 0.01. Functional annotation of the differentially expressed genes was performed using a web-based tool g:Profiler (Raudvere et al., 2019).

### Reverse transcription (RT) and quantitative qRT-PCR analysis

For RT-qPCR assays, CD11b+/F4/80+/Lyve1+ (Lp) and CD11b+/F4/80+/Lyve1- (Ln) macrophages were isolated by FACS. Sorted Lp and Ln macrophages were lysed with lysis buffer (RNeasy Mini Kit, Qiagen, Cat# 74104) and stored at -80°C. RNAs were then isolated with a QIAcube (Qiagen) instrument according to the manufacturer’s protocol with on column for RNase-Free DNase I (Qiagen, Cat# 79254) treatment. The quality of isolated RNAs was checked on the Agilent TapeStation with RNA Screen Tape (Agilent, Cat# 5067-5576). Reverse transcription was performed by synthesizing cDNAs from the Lp and Ln mRNAs with the Superscript III First Strand Synthesis System (Invitrogen, Cat# 18080051). RT-qPCR amplification was carried out using ready-to-use Taqman probe/primer sets (Applied Biosystems) to detect expression levels for *Lyve1* (*Mm00475056_m1*), *Folr2* (*Mm00433357_m1*), *Cd209f* (*Mm00471855_m1*), *Fcna* (*Mm00484287_m1*), *Timd4* (*Mm00724713_m1*), *Cd206* (*Mm01329362_m1*), *Igf1* (*Mm00439560_m1*), *Ang* (*Mm01316661_m1*), *Il1b* (*Mm00434228_m1*) and *Gapdh* (*Mm99999915_g1*) mRNAs. Two biological replicates (n=2 per replicate) were used for the Lp and Ln macrophages and assayed in triplicate. Relative RNA levels were calculated after normalizing to *Gapdh* mRNA using the 2^-ΔΔCt^ method, and the data were analyzed for significance using Student’s t-test.

### Flow cytometry-based phagocytosis assays

Macrophages were isolated from the hindlimb muscles of C57BL/6JN male mice as described above. Mononuclear cells from three animals were pooled for each set of experiment and cells were aliquoted for necessary treatment conditions and technical replicates. Three biological replicates (used total 9 mice) were analyzed. The phagocytic activity of macrophages was measured by red fluorescence from pHrodo E. coli bioparticles (Invitrogen, Cat# P35361). Briefly, 6 × 10^6^ macrophages were resuspended in 200 μL of Hams F10 complete media (Lonza, 12-618F) containing 10% horse serum (Gibco, 16050114) for each sample. Aliquots of 20 μL of pHrodo E. coli bioparticles, resuspended in live-cell imaging buffer (1 mg/ml, Invitrogen, Cat# A14291DJ) and sonicated for 2 min x 3, with 2 min intervals on ice between each sonication, were added to each cell tube, including appropriate FMO (fluorescence minus one) control tubes. Cell suspensions were gently and thoroughly mixed to ensure a homogenous distribution of the *E. coli* bioparticles. One set of samples was immediately transferred to a CO_2_ incubator for 2 h at 37°C, and another set (negative control) was incubated on ice for 2 h. After incubation, cells were washed with Live cell imaging solution at 400 rcf for 5 min, followed by another wash with PBS. All the steps were performed in the dark.

After the phagocytosis assay, cells were stained with viability dye followed by primary antibody staining as described above. Fluorochrome-conjugated antibodies used for staining the cells are as follows: BUV395 Rat Anti-Mouse CD45, PE-Cyanine7 anti-mouse/human CD11b Antibody, BUV737 Rat Anti-Mouse F4/80, Brilliant Violet 711™ anti-mouse I-A/I-E Antibody, APC Rat Anti-Mouse Lyve1 Antibody. The cells were acquired on a BD FACSAria™ Fusion instrument on the same day and analyzed with Flowjo. The relative phagocytosis levels were calculated using geometric Mean Fluorescence Intensity (gMFI), and the data were analyzed for significance using Student’s t-test.

### Immunofluorescent staining of macrophages in mouse SKM

Frozen sections from Rectus Femoris muscle from 3 m.o. C57BL/6J mice were cut, fixed in cold acetone, and subjected to regular double immunofluorescent staining or double TSA staining (Tyramide SuperBoost kit, Thermo Fisher, Cat# B40932) as performed previously (Cui et al., 2019). Primary antibodies recognizing Lyve1 (Abcam, Cat# ab14917, 1:200 dilution) and MHCII (Invitrogen, Cat# 14-5321-82, 1:100) worked well for regular immunostaining. Secondary antibodies were used for Lyve1 (Invitrogen, Cat# A-11012, 1:1000 dilution) and MHCII staining (Invitrogen, Cat# A-11006, 1:1000 dilution). Detection of CD11b (Santa Cruz, Cat# sc-1186, 1:50 dilution) required TSA staining. To identify LpHh macrophages in SKM, we carried out three sets of double staining: Lyve1 with MHCII by regular immunofluorescence staining, CD11b with Lyve1 and CD11b with MHCII by TSA staining. Photographs were taken using a DeltaVision microscope using a 20x lens.

## Author contributions

Linda K. Krasniewski, Investigation, Data curation, Formal analysis, Visualization; Papiya Chakraborty, Investigation, Data curation, Formal analysis, Visualization; Chang-Yi Cui, Conceptualization, Data curation, Formal analysis, Investigation, Visualization, Supervision, Writing – original draft, Writing – review and editing; Krystyna Mazan-Mamczarz, Data curation, Formal analysis, Visualization; Christopher Dunn, Investigation, Data curation, Formal analysis; Yulan Piao, Investigation, Data curation; Jinshui Fan, Investigation, Data curation; Changyou Shi, Investigation, Data curation; Tonya Wallace, Investigation, Data curation; Cuong Nguyen, Investigation, Data curation; Isabelle A. Rathbun, Data curation, Formal analysis; Rachel Munk, Investigation, Data curation; Dimitrios Tsitsipatis, Investigation, Data curation, Formal analysis; Supriyo De, Data curation, Formal analysis; Payel Sen, Investigation, Data curation; Luigi Ferrucci, Conceptualization, Supervision; Myriam Gorospe, Conceptualization, Supervision, Writing – review and editing.

## Acknowledgements

This work was supported by the Intramural Research Program of the National Institute on Aging. The authors thank Marc Michel for technical assistance.

## Data availability statement

The single-cell RNA-seq analysis was uploaded to GEO with identifier GSE195507; token for reviewers is etifyqaexrcrjmh.

## LEGENDS FOR SUPPLEMENTARY FIGURES

**Fig. S1:**
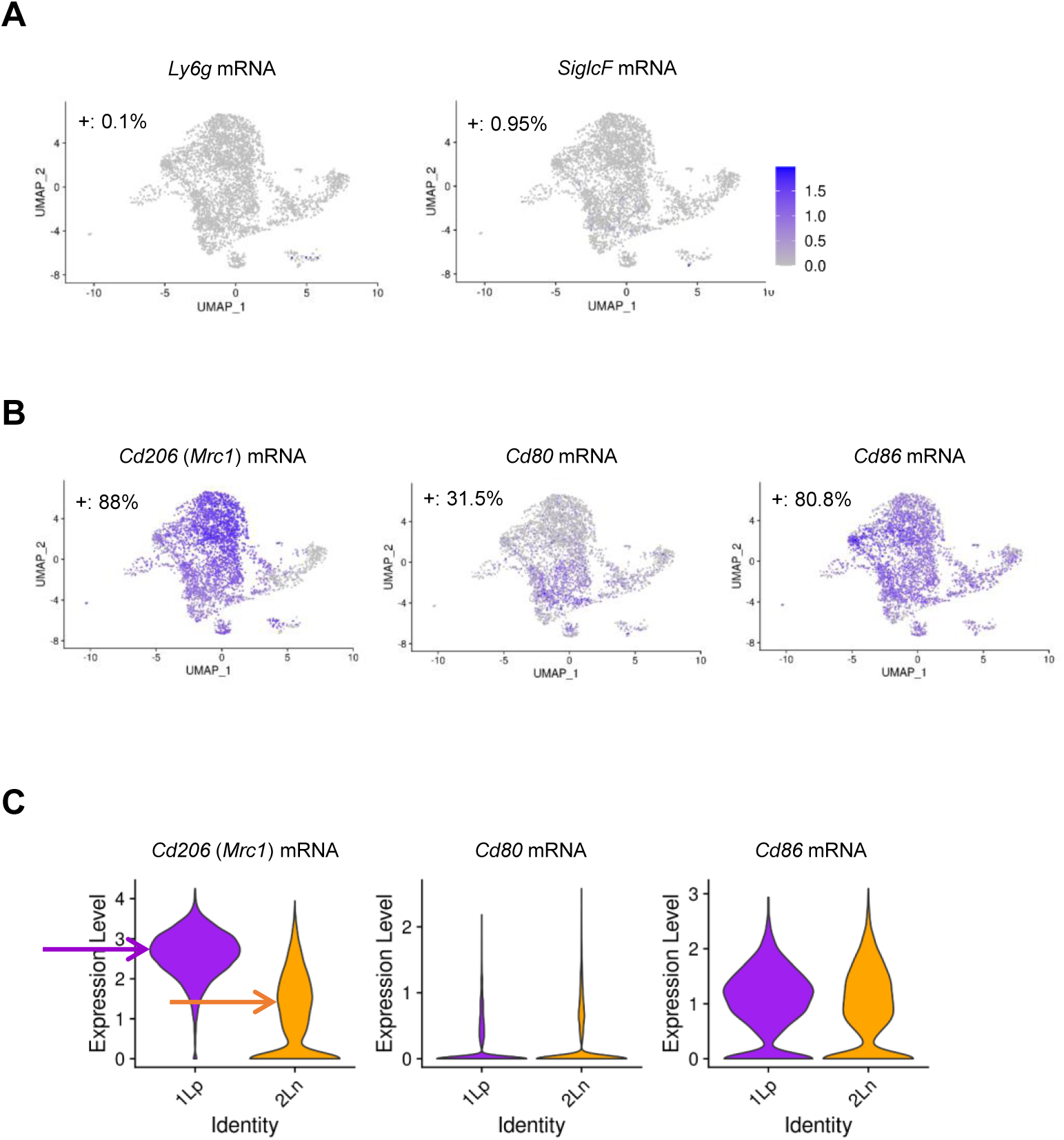
Quality control experiments for the SKM macrophage, single-cell RNA-seq analysis. (A) UMAP analysis of minimal contamination of neutrophils and eosinophils in the CD11b+/F4/80+ macrophage population. (B) *Cd206* and *Cd86* mRNAs, widely expressed in >80% of total macrophages; most macrophages expressed both *Cd206* and *Cd86* mRNAs, while *Cd80* mRNA was expressed in a small population. (C) Expression levels of *Cd206, Cd80* and *Cd86* mRNAs in LYVE1+ and LYVE1-macrophages. *Cd206* and *Cd86* mRNAs were highly expressed in LYVE1+ macrophages, but also expressed in LYVE1-macrophages. Both LYVE1+ and LYVE1-macrophages expressed low levels of *Cd80* mRNA. Arrows point to mean expression levels of *Cd206* in Lp and Ln macrophages.

**Fig. S2:**
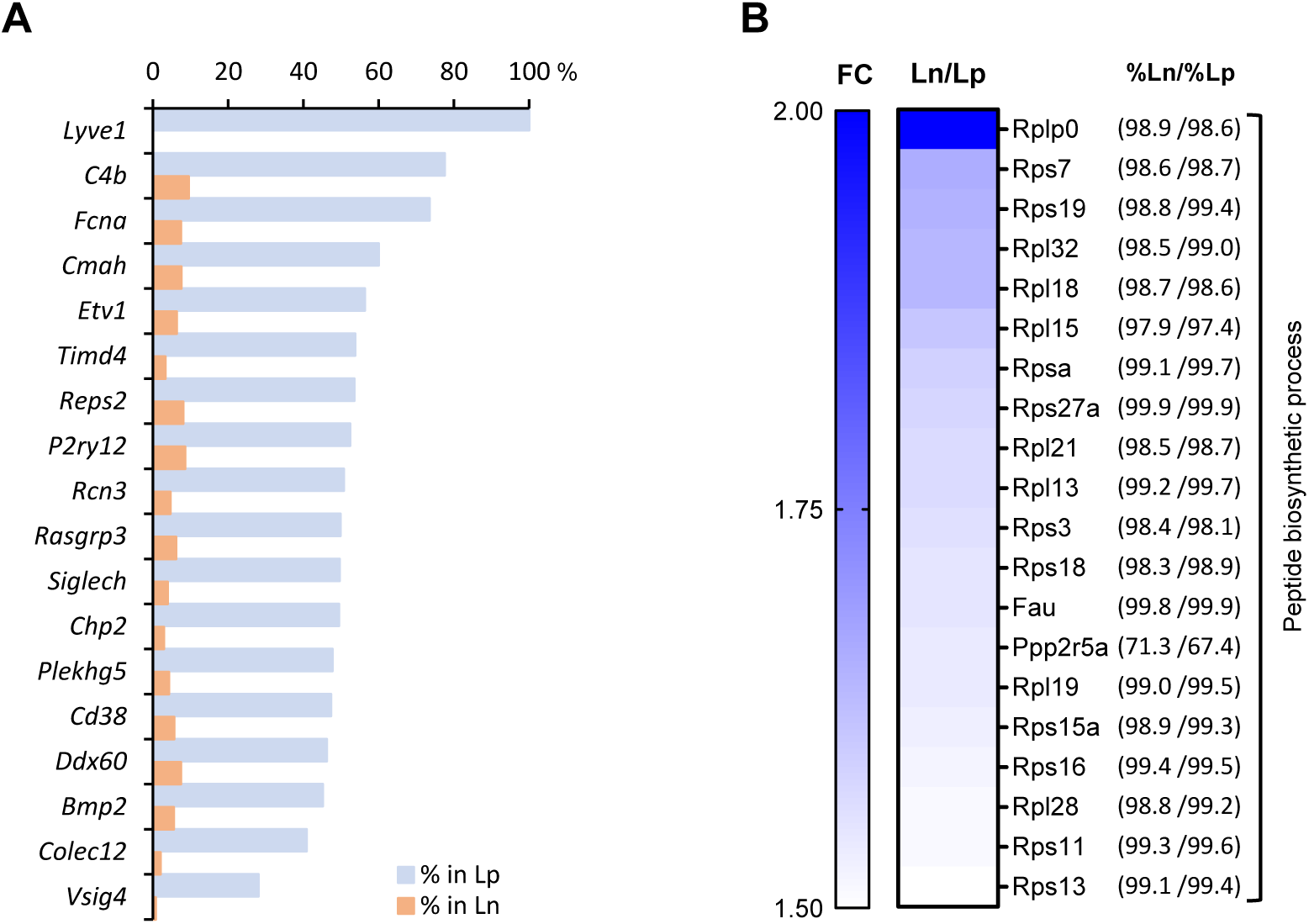
mRNAs highly expressed in LYVE1+ or LYVE1-macrophages. (A) mRNAs almost exclusively expressed in LYVE1+ (Lp) population. (B) Cluster of mRNAs encoding ribosomal proteins that was highly expressed in LYVE1- (Ln) macrophages. Note, genes listed in this panel were expressed in similar number of macrophages in the Lp and Ln subpopulations (%Lp/%Ln), but were highly expressed in the Ln subpopulation (Ln/Lp).

**Fig. S3:**
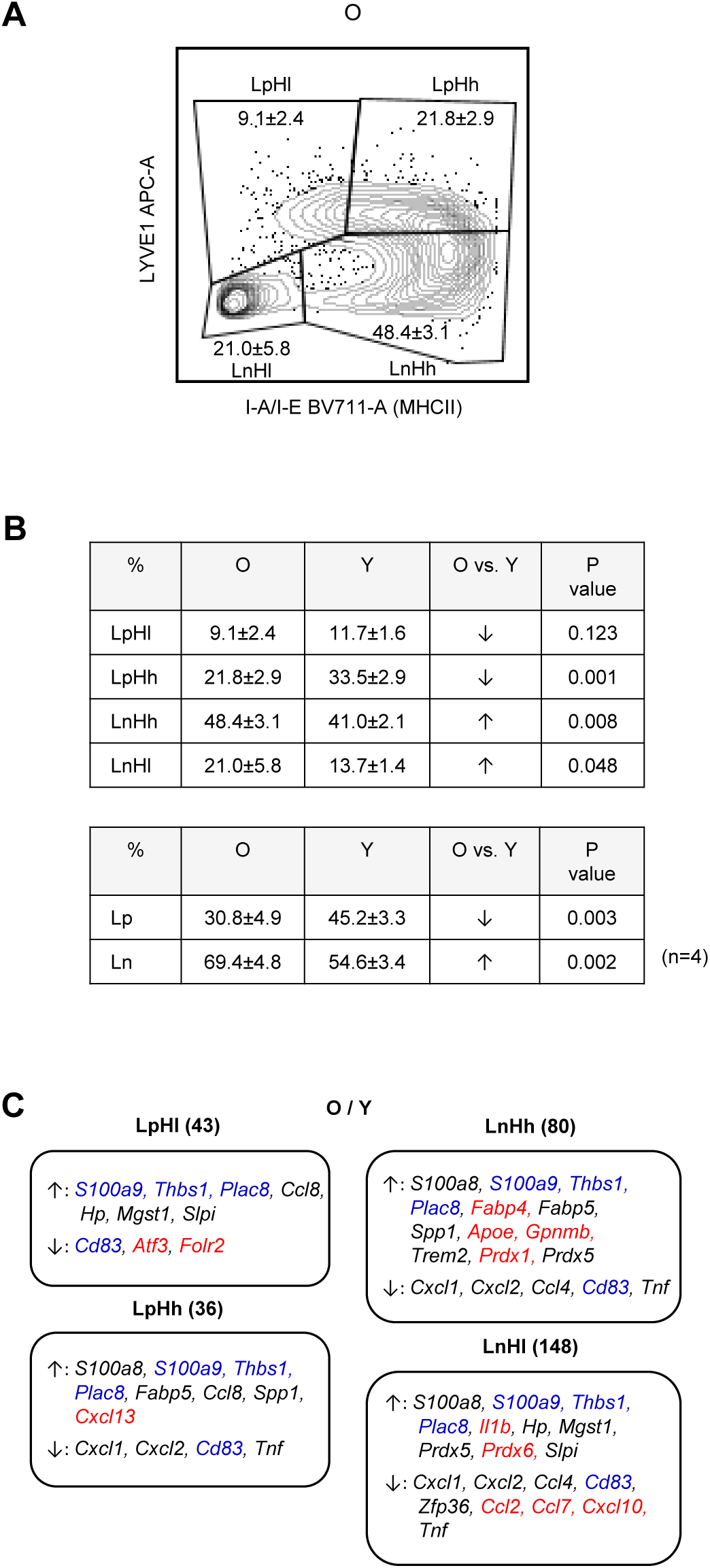
Changes in genes expressed by macrophages from old SKM. (A) Flow cytometric analysis of macrophages from old SKM showing four subgroups, as in the young (main Fig. 3e). Gating was based on FMO controls for each experiment. (B) In old SKM (relative to young), the number of macrophages in the LpHl and LpHh subgroups decreased, while the number of macrophages in the LnHh and LnHl subgroups increased. *Top*, reduced LpHh and increased LnHh in old relative to young SKM macrophages. *Bottom*, reduced Lp and increased Ln in two-group classification by LYVE1 marker only. (C) Select mRNAs differentially expressed in young and old SKM macrophage subgroups. Blue, mRNAs significantly changed in all four subgroups. Red, mRNAs significantly altered in one subgroup only. Numbers in parentheses indicate the number of genes differentially expressed in young and aged SKM macrophages in each subgroup.

## Supplementary tables

**Table S1:** mRNAs highly expressed in LYVE1+ or LYVE1-macrophages

**Table S2:** mRNAs in each of four macrophage subgroups

**Table S3:** Differentially expressed mRNAs among subgroups

**Table S4:** Differentially expressed mRNAs in total and each subgroup of macrophages from SKM from young and old mice

**Table S5:** Featured mRNAs in unsupervised clusters and abundance changes in old mice

